# Clade-wide fungal proteome analysis reveals structure–function conservation in divergent Dicer proteins

**DOI:** 10.1101/2025.04.28.651110

**Authors:** Lorena Melet, Jonathan Canan, Pablo Villalobos, Boris Vidal-Veuthey, Fabian González-Toro, Ivana Orellana, J. Andrés Rivas-Pardo, Juan P. Cárdenas, Carol Moraga, Víctor Castro-Fernandez, Nathan R. Johnson, Elena A. Vidal

## Abstract

Dicers (Dcrs) are central proteins involved in the biogenesis of small RNAs (sRNAs) in eukaryotes. Most of the knowledge on Dcr structure, function and evolution comes from studies conducted in animal and plant species. Comparatively, much less is known in fungi, which are a genetically and ecologically diverse group with important roles in ecosystems, agriculture, medicine, and biotechnology. While canonical Dcrs in plants and animals contain a well-defined domain architecture, most fungal Dcrs with experimentally validated functions lack one or more identifiable canonical domains, raising questions about how RNA-binding and precise sRNA processing is retained. Here, we conducted the most extensive survey of fungal Dcr proteins, analyzing 1,593 proteomes across eight phyla. We found a diversity of Dcr domain architectures, with some of them lacking an identifiable PAZ, Helicase, and/or double-stranded RNA binding domains. Phylogenetic analyses showed that different Dcr classes are distributed across distinct clades that often align with fungal taxonomic groups. Despite the lack of canonical domain architectures, we found that fungal Dcrs fold into a characteristic L-shaped structure and show PAZ-like folds, even in proteins without detectable PAZ sequences. Molecular docking and electrostatic analyses further indicate that these divergent Dcrs maintain key RNA-binding surfaces for proper sRNA processing. Our results indicate a remarkable evolutionary plasticity of Dcr in fungi, showing that essential sRNA processing functions can be retained through structural conservation, and highlighting fungi as models to study the modular evolution of the RNAi machinery in eukaryotes.

**Significance statement:** Dicer (Dcr) proteins are central to RNA interference (RNAi), a gene regulatory mechanism conserved across eukaryotes. However, current models of Dcr structure, function, and evolution are largely based on studies in animals and plants. Here, we present the most comprehensive analysis to date of Dcr proteins in fungi, a diverse eukaryotic group including many societally important pathogens and symbiotes which are reliant on RNAi. Our findings reveal that despite widespread divergence from canonical Dcr architecture, fungal Dcrs conserve critical folds and RNA-binding features, further suggesting that core RNAi functions are maintained. This work establishes fungi as key models for studying the evolution and functional robustness of the RNAi machinery, offering broader insight into the diversity and plasticity of sRNA biogenesis pathways across eukaryotes.

## Introduction

RNA interference (RNAi) is a conserved eukaryotic mechanism with diverse roles, including gene expression regulation (1), antiviral defense (2), genome stability (3), and communication between cells and across species (4–6). RNAi is mediated by small RNA (sRNA) molecules broadly categorized as 18-30 nucleotides in length, which are generated through the cleavage of exogenous or endogenous double-stranded RNA (dsRNA) precursors. These sRNAs are loaded into RNA-induced silencing complexes, which use sequence complementarity to target nucleic acid sequences for transcriptional or post-transcriptional gene silencing (7, 8). Among the proteins responsible for processing sRNAs from dsRNAs, Dicer (Dcr) plays a central role, particularly in the small interfering RNA (siRNA) and microRNA (miRNA) pathways.

Numerous studies have elucidated the functions of Dcr domains in recognizing and processing their canonical substrates, including double-stranded RNAs and single-stranded hairpin precursors of miRNAs (pre-miRNAs). Structurally, Dcr proteins exhibit an L-shaped, modular, multidomain architecture that can be broadly divided into three regions along their three-dimensional organization: a base, a core (body), and a head. These regions reflect both domain composition and spatial-functional arrangement (9–11). The base (N-terminal region) contains the helicase domain, comprised of three subdomains: DExD/H-box, RESIII, and helicase C-terminal domains. These serve as the entry point for dsRNA substrates, recognizing and binding them before unwinding the RNA duplex to ensure proper positioning. Evidence shows that in some species and Dcr variants the helicase domain may or may not be involved in ATP hydrolysis, a function that has been related to recognition and processing of blunt-ended dsRNAs such as viral RNAs or endogenous dsRNAs (12–14). The core (central region) includes the domain of unknown function (DUF283) and double-stranded RNA binding domain (dsRBD). DUF283 functions primarily as a protein–protein interaction module, selectively interacting with dsRBD-containing cofactors such as DRB4 and HYL1 in Arabidopsis (15, 16), or proteins such as ADAR1 in mammals (17). The dsRBD enhances the stability and specificity of enzyme-substrate interaction contributing to the fidelity of the cleavage reaction (18). At the catalytic core, the tandem RNase IIIa and IIIb domains form a dimer that cleaves both strands of the dsRNA duplex. The head (C-terminal region) contains the Piwi/Argonaute/Zwille (PAZ) domain, which binds the 3’ overhangs of dsRNA substrates, acting as an anchoring point that guides the cleavage site (19). In the three-dimensional structure of Dcr, the PAZ domain is connected to the RNase III catalytic center by a connector α-helix. This helix acts as a molecular ruler, setting the distance between the RNA-binding PAZ domain and the RNase III domains to define the length of the sRNAs produced (∼21–25 nt) (20).

While this canonical domain architecture is conserved across most of the Dcr proteins in animals and plants, reduced Dcr proteins have also been described. Remarkably, the Dcr protein from the protist *Giardia intestinalis* consists solely of the tandem RNase III domains connected through a connector helix to a platform region, yet remains fully competent in cleaving dsRNA substrates (20). Interestingly, many Dcr proteins in fungi have been reported to lack identifiable Helicase, PAZ and/or dsRBD domains, suggesting a widespread presence of non-canonical Dcrs (21–23). This diversity positions fungal Dcrs as a valuable model for exploring how ancestral RNase III enzymes evolved into the complex, multidomain Dcr proteins found in higher eukaryotes.

Fungi comprise a large and diverse group of eukaryotic organisms, currently classified into at least 12 major phyla (24), with some recent taxonomic studies recognizing up to 18 or 19 distinct phyla based on phylogenomic evidence (25–27). Most described fungal species to date belong to the higher fungi Ascomycota and Basidiomycota, which together form the clade Dikarya. Other fungal phyla (e.g., Mucoromycota, Chytridiomycota, Zoopagomycota) represent a smaller portion of known fungal diversity, though they are ecologically important and likely under-sampled (25).

Multiple approaches have been used to identify RNAi-related proteins in fungi, ranging from focused studies in groups of species (22, 23, 28) to large-scale comparative surveys such as the FunRNA database, which catalogued RNAi components across 131 fungal genomes (21). These studies have shown that the number and architecture of Dcr proteins vary significantly across fungal species. While many fungi encode two Dcr paralogs (29, 30), these can differ in sequence and domain organization, which may reflect functional specialization (22, 31). However, there are also fungal lineages where only a single Dcr has been identified (22, 32) and others, including members of the Saccharomycotina and some Basidiomycota, where Dcr genes appear to be absent altogether (21–23). Notably, the apparent absence of Dcr domains, such as PAZ or dsRBD in some fungal proteins may result from limitations in domain annotation pipelines rather than true biological loss. For example, domains like PAZ may be highly divergent in fungi, and thus escape identification by current probabilistic models based on protein alignments of animal and plant-derived PAZ domains (33). However, albeit divergent, PAZ-like folds have been reported in fungal Dcrs such as Dicer from *S. pombe* (34), hinting at tridimensional structure conservation, and at the importance of incorporating structural analyses to generate hypotheses on fungal Dcr function.

The number of publicly available annotated and reference genomes has expanded more than tenfold in the last years, presenting a prime opportunity to study Dcr diversity at a wide phylogenetic scale. Furthermore, integration of structure prediction methods like AlphaFold (35) can help us determine whether structural conservation is widespread across fungal Dcrs despite apparent sequence diversity.

In this study, we present a comprehensive analysis of Dcr proteins across fungi, revealing widespread non-canonical architectures that often lack identifiable PAZ, Helicase, or dsRBD domains. Through a combination of phylogenetic analyses, structural modeling, and molecular docking predictions, we show that many of these divergent Dcrs retain key features for RNA processing, including structurally conserved PAZ-like regions. Our findings expand the understanding of Dcr diversity in fungi and suggest that RNAi functionality can be maintained despite extensive domain loss or divergence.

## Results

### Comprehensive identification of Dcr proteins in fungi

To comprehensively catalog fungal Dcr proteins, we retrieved all available fungal reference proteomes from the NCBI, obtaining 1,593 datasets representing 1,427 unique species (Supplementary Table 1). The selected proteomes were predicted from genomes labeled as ’reference’ or ’annotated’ in NCBI, excluding those classified as atypical. These genomes meet the criteria for reference-quality assemblies. Most of the selected proteomes exhibit high completeness, with 86.4% scoring over 90% in BUSCO completeness based on either the fungi-wide or phylum-specific datasets (Fig. 1A, Supplementary Table 1). The proteomes span a wide range of fungal organisms with a marked representation from the phyla Ascomycota (913 species, 64.1%) and Basidiomycota (333 species, 23.4%). However, non-Dikarya fungi also constitute a significant proportion of the proteomes, with over 172 species from Mucoromycota, Zoopagomycota, Chytridiomycota, Microsporidia, Blastocladiomycota, and Olpidiomycota (Supplementary Table 1). This dataset expands the taxonomic scope for the identification and comparative analysis of fungal Dcr proteins, as compared with previous studies (21, 23, 28).

**Figure 1.**
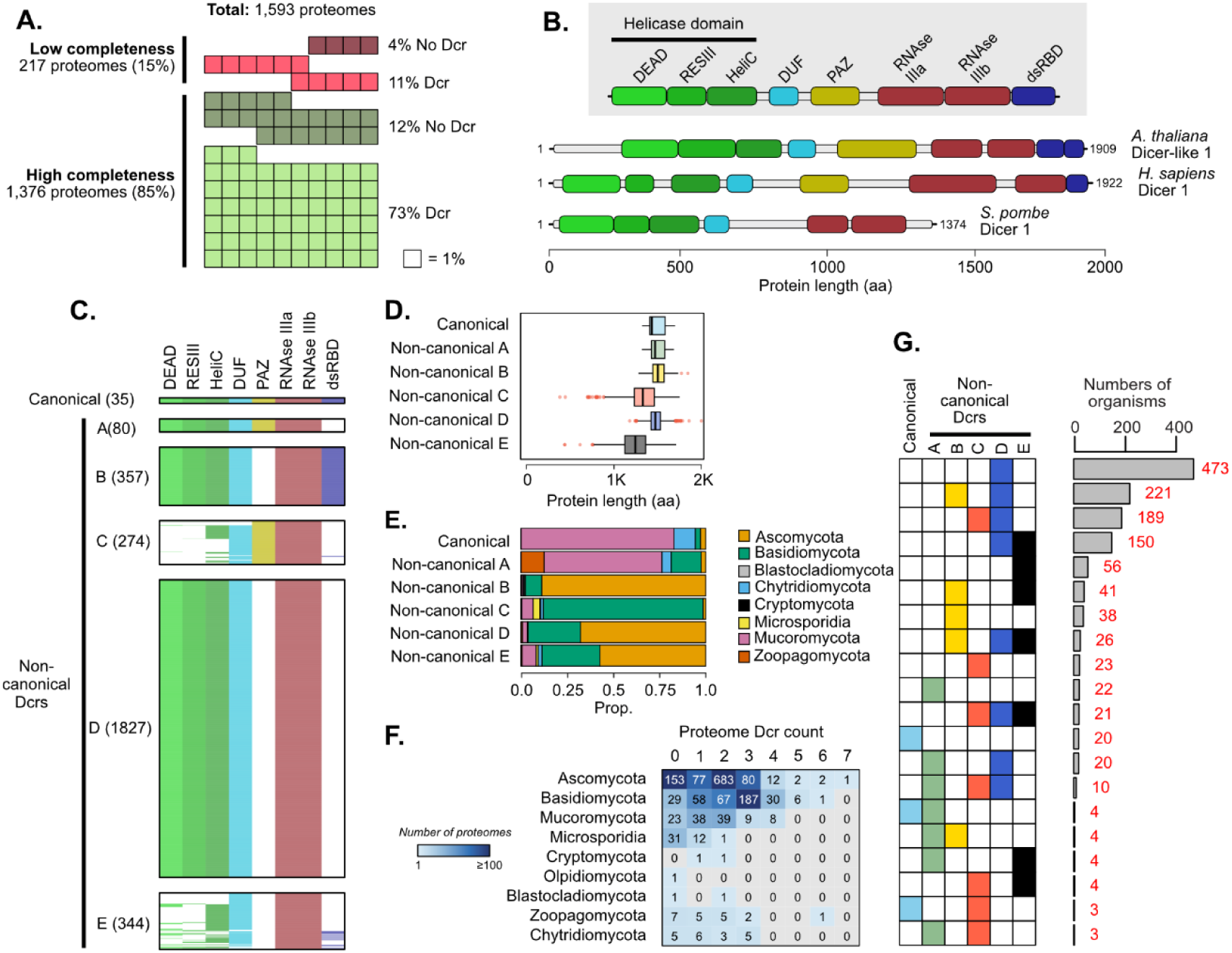
Domain-based classification and distribution of Dcr proteins across fungal phyla. (**a**) Summary of the 1,593 annotated fungal proteomes analyzed for the identification of Dcr proteins, showing the percentage of low (<90%) and high (>90%) completeness proteomes according to BUSCO. (**b**) Representative domain architectures of known Dcr proteins from *Arabidopsis thaliana*, *Homo sapiens*, and *Schizosaccharomyces pombe*, illustrating the conserved domain composition: DEAD, ResIII, Helicase C (HeliC), DUF283, PAZ, RNase IIIa and IIIb, and dsRNA-binding domain (dsRBD). Protein lengths are shown in amino acids (aa). (**c**) Classification of all predicted fungal Dcr proteins into six structural categories based on the presence or absence of key domains. Canonical Dcrs retain a full complement of conserved domains, while non-canonical groups exhibit partial or divergent architectures. (**d**) Average protein length (in aa) per Dcr category. (**e**) Distribution of Dcr categories across eight fungal phyla, highlighting differences in domain conservation and structural diversity among lineages. (**f**) Frequency distribution of fungal species harboring one to seven Dcr copies, indicating variability in Dcr gene family expansion across the dataset. (**g**) Combinations of Dcr categories across fungal genomes. Each row shows a unique combination of canonical and non-canonical Dcrs (A–E), with colored squares indicating presence of a certain domain. Bars on the right represent the number of organisms per combination (in red).

To identify putative Dcr proteins, we screened the proteomes for canonical Dcr-associated domains previously described in plants and animals. Domain searches were performed using HMMER, following methodologies established for Dcr identification in fungi (21) and other organisms (36). A protein was classified as a putative Dcr if it met two criteria: (1) the presence of two, tandem RNase III domains, essential for dsRNA cleavage, and (2) at least one RNA-binding domain, including Helicase subdomains DExD/H-box, ResIII or HeliC, DUF283, PAZ, or dsRBD. This approach identified 2,917 putative Dcr proteins across 1,215 species from eight fungal phyla (Supplementary Table 2). To validate these predictions, we used an independent domain annotation approach using InterProScan, which showed 100% concordance with the HMMER-based results (Supplementary Table 2).

Interestingly, we found no evidence of Dcr proteins in 12% of BUSCO high-completeness proteomes and 11% of BUSCO low-completeness proteomes, totaling 227 species (254 proteomes). This suggests that Dcr proteins may have been lost in up to 15% of the surveyed fungal species. However, in proteomes with low completeness scores the absence of Dcr should be interpreted with caution, as it may reflect incomplete gene models rather than true gene loss. Among the proteomes with no detectable Dcrs we identified 127 Ascomycota species, of which 115 (91%) belong to the subphylum Saccharomycotina, including members from the classes Dipodascomycetes, Saccharomycetes, Pichiomycetes and Trigonopsidomycetes (Supplementary Table 3). This subphylum encompasses most Ascomycete yeasts, several of which have previously been reported to lack a functional RNAi machinery (22). In addition, species lacking Dcrs were also found in Basidiomycota, Olpidiomycota, Blastocladiomycota, Chytridiomycota, Microsporidia, Mucoromycota, and Zoopagomycota (Supplementary Table 3), suggesting that Dcr protein loss has occurred multiple times independently across fungal lineages.

### Widespread domain architecture variation and predominance of non-canonical Dcr types in fungi

To characterize the domain composition of fungal Dcrs, we referred to the canonical domains typically found in Dcr proteins (Fig. 1B). Using these domain features as a guide, we classified the 2,917 identified Dcr proteins into six categories based on the presence or absence of domains detected through HMMER analysis. The types of domains found across the fungal dataset is summarized in Fig. 1C, revealing the predominance of non-canonical forms and the high frequency of Dcr proteins where the PAZ and/or dsRBD domains could not be detected. Proteins containing all expected domains were classified as canonical Dcrs, representing a small subset of the dataset (35 proteins, 1.2%). Proteins with one or more missing domains were designated as non-canonical Dcrs and further grouped according to the specific patterns of domain loss. Non-canonical A includes proteins lacking only the dsRBD domain (80 proteins, 2.7%), while non-canonical B consists of those missing the PAZ domain but retaining all other domains (357 proteins, 12.2%). Non-canonical C comprises proteins in which Helicase subdomains and dsRBD are absent (274 proteins, 9.4%). The most abundant group, non-canonical D, consists of proteins lacking both the PAZ and dsRBD domains (1,827 proteins, 62.6%). Finally, non-canonical E includes proteins in which three or more domains could not be detected (344 proteins, 11.8%).

The PAZ domain, central to RNA binding and precise determination of sRNA length, was undetectable in the vast majority of Dcr proteins, with 86.7% lacking a recognizable PAZ domain. However, protein length analysis revealed that proteins without a detectable PAZ, particularly those in the non-canonical B and D categories, are comparable in size to canonical and non-canonical A proteins that do contain this domain (Fig. 1D). This suggests that the PAZ domain may still be present but has undergone substantial sequence divergence relative to the canonical PAZ (cPAZ) described in plants and animals, rendering it undetectable by standard HMM-based methods. Consistent with this idea, the current PAZ HMM matrix used in this analysis (PF02170/IPR3100) was constructed from Dcrs and ARGONAUTE proteins mainly from plants and animals, with negligible representation of fungal Dcrs. Similarly, the dsRBD domain, also involved in RNA binding, was frequently undetected, with 84.7% of proteins missing this domain. Interestingly, proteins in the non-canonical C and E categories, which often lack subdomains of the Helicase region as well as the dsRBD domain, exhibited shorter lengths, suggesting that the absence of detectable domains in these groups may in some cases result from protein truncation rather than sequence divergence.

The distribution of Dcr categories varies substantially among fungal phyla (Fig. 1E). Canonical Dcrs and non-canonical A proteins, both of which retain a cPAZ domain, are enriched in Mucoromycota, a phylum composed of mychorrizal symbionts, plant and animal pathogens, fungal parasites and decomposers of organic matter (37). In contrast, non-canonical C proteins, which also retain a cPAZ domain, were primarily found in Basidiomycota. Non-canonical B, D, and E proteins, lacking a cPAZ domain, are most prevalent in Ascomycota, with some also found in Basidiomycota. The number of dcr genes per genome also varied by phylum; most Ascomycota species have two dcr genes (683 species, 68%) while Basidiomycota more frequently possess three (187 species, 49.5%). Strikingly, Microsporidia, a group of obligate intracellular parasites, either lacked dcrs entirely (31 species, 70.5%) or contained only a single dcr (12 species, 27.3%) (Fig. 1F), consistent with previously described reductions in gene contents in this lineage (38).

We also analyzed the distribution of Dcr category combinations across fungal species (Fig. 1G). Most species contain non-canonical D Dcrs as the sole Dcr type (473 species), or in combination with non-canonical B (221 species), C (189 species), or E (150 species). Interestingly, canonical Dcrs were either found alone or in combination with non-canonical A or C, both of which retain a cPAZ domain. These patterns underscore the diversity in both the number and types of Dcr proteins across fungal taxa, suggesting lineage-specific adaptations in the RNAi machinery, likely in response to differing ecological and/or evolutionary pressures.

### Clade-specific distribution of canonical and non-canonical Dcrs suggests independent diversification of RNAi pathways in fungi

To investigate the phylogenetic relationships of fungal Dcr proteins, we constructed a sequence similarity network, considering a cut-off of 80% coverage and 50% identity between connected nodes. 2,636 Dcr proteins met these criteria and were clustered into 149 groups (Supplementary Table 2). Multiple sequence alignments were generated for each cluster and integrated into a global alignment, which was used to construct a maximum-likelihood phylogenetic tree, including outgroup sequences from plants and metazoans (Fig. 2A). The analysis revealed a set of well-supported clades that we defined as major Dcr groups. These groups correspond to fungal taxonomic lineages, with Ascomycota sequences separating into two major clades (As1 and As2) and Basidiomycota sequences into three distinct clades (Ba1, Ba2, Ba3). In contrast, sequences from non-Dikarya fungi formed more scattered groups, forming independent clades (Mu1, Mu2, Mi1, Bl1, Ch1), suggesting that Dcr diversification in non-Dikarya followed an independent evolutionary trajectory. The outgroup sequences formed a distinct branch, supporting the overall topology and robustness of the tree (Fig. 2A).

**Figure 2.**
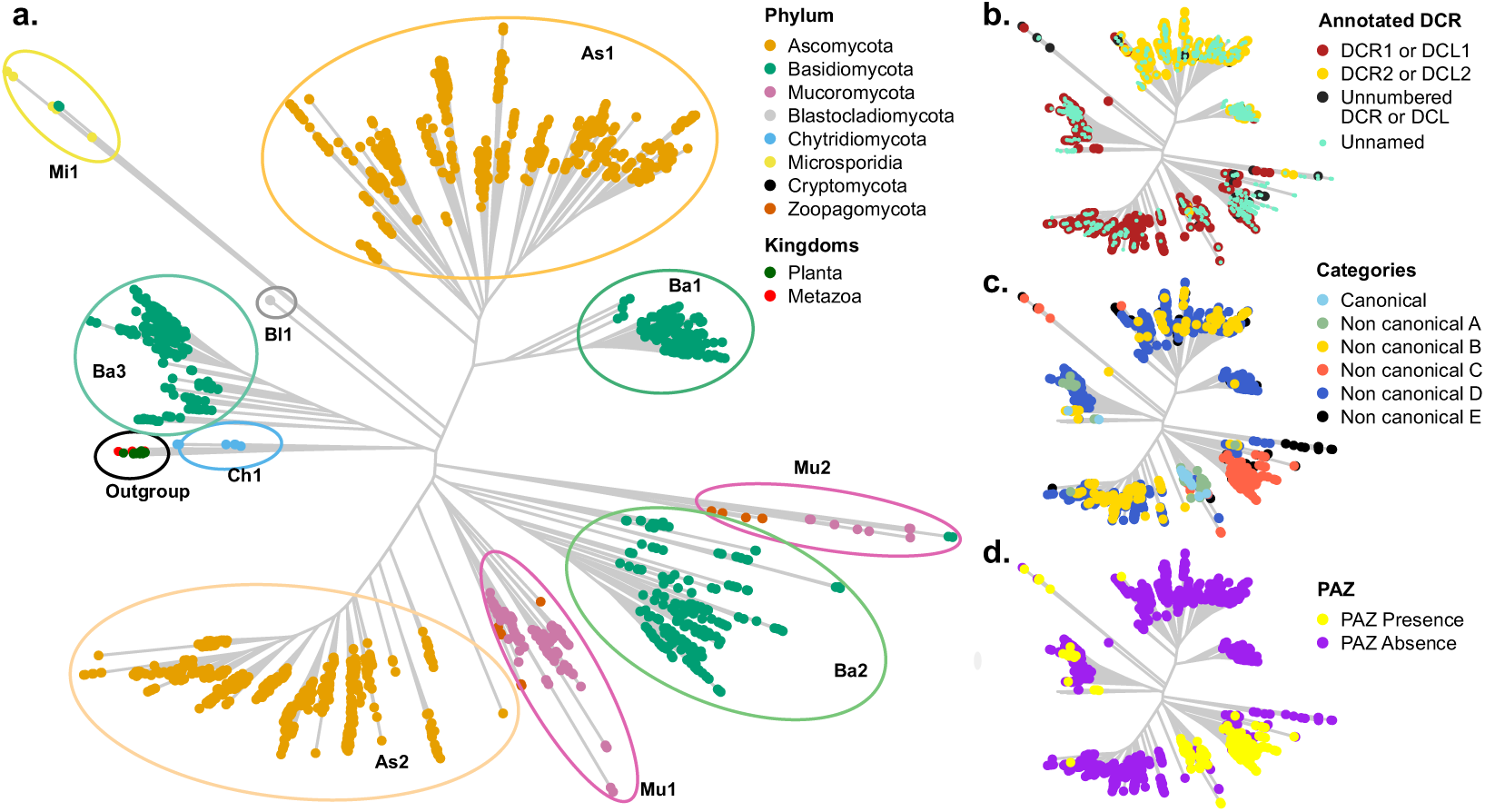
Phylogenetic and functional classification of fungal Dcr proteins. (**a**) Maximum-likelihood phylogenetic tree showing the distribution of Dcr proteins across eight fungal phyla, as well as representative outgroups from plants and metazoans. Colored ovals highlight major clades identified within fungal lineages, labeled as As1–As2 (Ascomycota), Ba1–Ba3 (Basidiomycota), Mu1–Mu2 (Mucoromycota and Zoopagomycota), and others, based on their phylogenetic position. (**b**) Annotation status of Dcr proteins according to UniProtKB, distinguishing Dcr1/Dcl1 (Dark red), Dcr2/Dcl2 (yellow), unnumbered Dcr/Dcl entries (black), and unnamed proteins (light green). (**c**) Classification of fungal Dcrs into canonical and five non-canonical categories (A–E), based on domain composition. (**d**) Mapping of cPAZ domain presence (yellow) or absence (purple) across the phylogeny.

To evaluate whether clade assignments relate to previously reported Dcr annotations, we mapped proteins annotated as Dcrs in UniProtKB onto the phylogenetic tree (Fig. 2B). These include proteins labeled as Dcr1, Dcr2, Dcr-like1 (Dcl1), Dcr-like2 (Dcl2), as well as others annotated more generically as “Dicer” or “Dicer-like”. We found that proteins annotated as Dcr1 or Dcl1 are predominantly located within the Ascomycota As2 clade and the Basidiomycota clades Ba2 and Ba3. Dcr1/Dcl1 proteins are also observed in the Mu1 and Mu2 clades, as well as some identified as Dcr2/Dcl2 in Mu2. These ancient duplications forming Ba2/3 and Mu1/2 point to an unclear resolution of the ancestral proteins and possible paraphyletic naming. However, these are the exception. In general, Dcrs with similar annotations tend to group within the same clades, suggesting that existing naming conventions reflect underlying evolutionary relationships, even if their functional roles have not been fully confirmed. Notably, most of the Dcrs identified in our analysis remain unannotated in public databases (Fig. 2B, Supplementary Table 2), underscoring the extensive under-representation of fungal Dcr diversity and abundance.

To further explore the relationship between phylogeny and domain architecture, we mapped canonical and non-canonical classifications onto the phylogenetic tree. Canonical Dcrs clustered primarily within the Mu1 clade, and to a lesser extent, in the Mu2 clade, consistent with their enrichment in the Mucoromycota phylum (Fig. 1E, Fig. 2C). Non-canonical A and C proteins, which, as canonical Dcrs, retain a cPAZ domain (Fig. 1C, Fig. 2D), also show a restricted distribution, with non-canonical C primarily associated with the Ba2 clade and non-canonical A with Ba3. In contrast, non-canonical B, D and E proteins, which lack a cPAZ domain (Fig. 2D), were widely distributed across the tree (Fig. 2C). The restricted phylogenetic distribution of cPAZ-containing Dcrs suggests this domain is not broadly conserved across fungal lineages; rather, its retention appears to be lineage-specific and may reflect convergent retention or loss events in different evolutionary contexts.

The helicase subdomains DExD/H-box, ResIII, and HeliC also showed differential representation across fungal clades, although most Dcrs retained Helicases with all three subdomains (Fig. 1C, Supplementary Fig. 1A). Notably, Dcrs lacking all helicase components, or retaining only the HeliC subdomain, were mainly found in the Ba2 clade, suggesting that Basidiomycota harbor both helicase-complete Dcrs (e.g., in Ba1 and Ba3) and more divergent forms potentially lacking ATPase activity (Ba2). A similar pattern was observed in Mucoromycota, with Mu1 Dcrs containing complete helicases and Mu2 Dcrs showing partial loss of subdomains. Additional incomplete helicase architectures were also found in the Microsporidian clade Mi1. In contrast, the dsRBD domain, another typical component of canonical Dcrs, but rarely detected in our analysis, was present in proteins found in diverse clades across the tree (Supplementary Fig. 1B). These results underscore the evolutionary plasticity of fungal Dcrs, suggesting that distinct selective pressures have driven domain loss, retention, or divergence across fungal lineages.

### Fungal Dcrs have conserved structural features despite sequence divergence

The sequence similarity network and phylogenetic analyses suggest a high degree of sequence divergence among fungal Dcr proteins, with several instances in which key domains typically associated with Dcr function were not detected, likely due to either their absence or low sequence conservation. To explore whether these proteins might nonetheless retain conserved three-dimensional features, we generated structural predictions for all Dcr proteins using AlphaFold (35). Protein models displaying a predicted Local Distance Difference Test (pLDDT) score greater than or equal to 70 were selected, and a structural similarity network was constructed, considering a template modeling (TM) score of ≥ 0.9 between connected nodes as a threshold. Among the selected models, the average pLDDT score was 75, supporting the overall reliability of the structural predictions. A total of 2,475 proteins met this criterion and were grouped into 26 clusters (Supplementary Table 2). Strikingly, nearly 95% of phylogenetically analyzed proteins are found within just four major structural clusters (Fig. 3A), suggesting core structural features are widely conserved. Each cluster showed a distinct representation of fungal phyla (Fig. 3A, B). Clusters 1 and 2 contain the majority of Dcr proteins, with cluster 1 including predominantly Ascomycota sequences and a smaller subset from Basidiomycota. This cluster harbors Dcrs with experimentally validated roles in sRNA processing such as *Botrytis cinerea* Dcr2 (6, 39, 40), *Aspergillus fumigatus* Dcr2 (41) or *Trichoderma atroviride* Dcr2 (42–44). Cluster 2 is predominantly composed of Ascomycota sequences, and includes *A. fumigatus* Dcr1, *T. atroviride* Dcr1 and *Neurospora crassa* Dcr1 (45). Clusters 3 and 4 include fewer proteins, mainly from Basidiomycota and Mucoromycota (cluster 3) or exclusively from Basidiomycota (cluster 4). The remaining smaller clusters consist of Ascomycota and Basidiomycota Dcrs and may represent functionally divergent paralogs present in these phyla (Fig. 3A, B).

**Figure 3.**
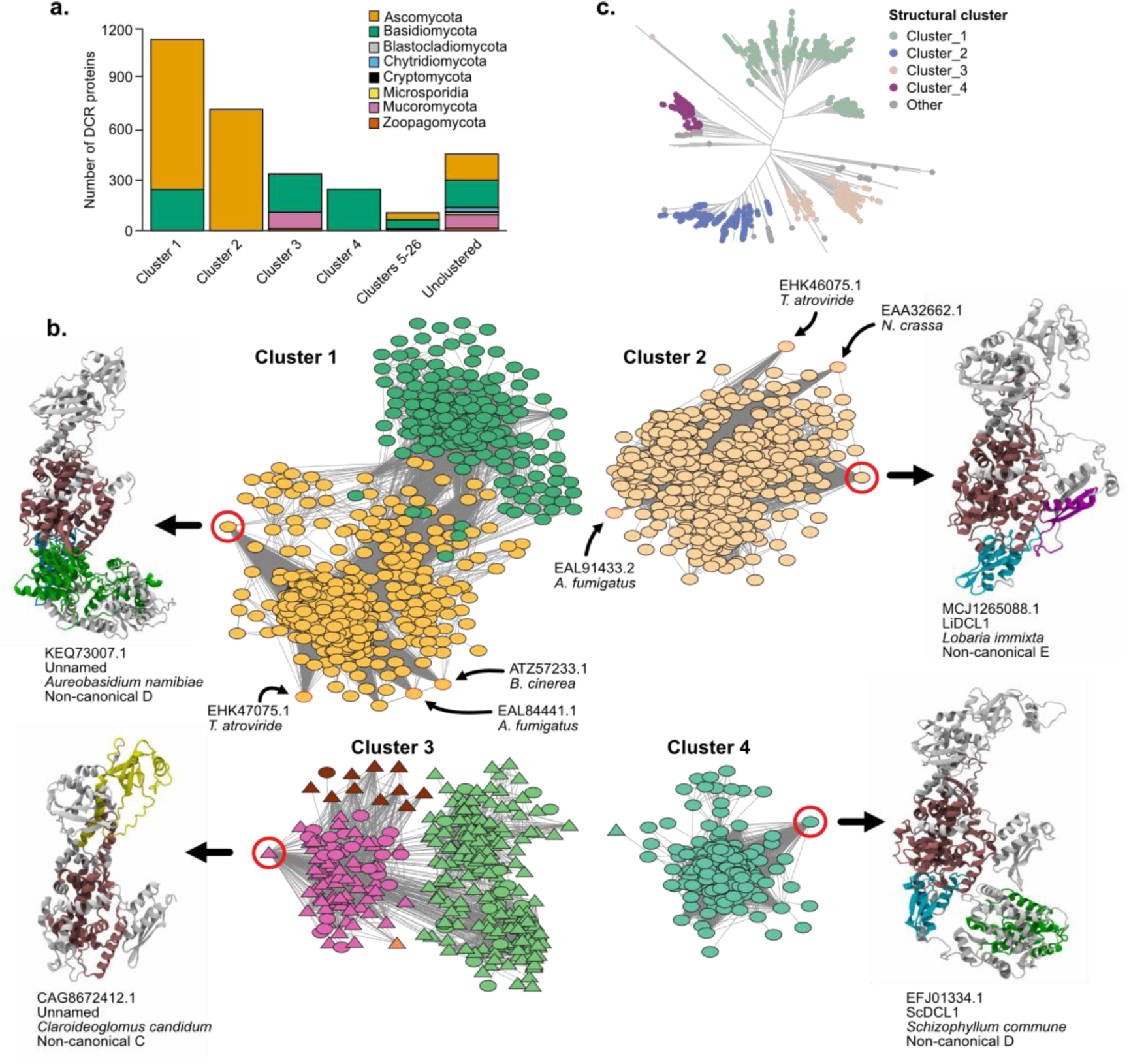
Structural Network of Proteins Revealing Functional Clusters Based on Folding Patterns. (**a**) Taxonomic composition of each structural cluster, shown as the number of proteins per fungal phylum. (**b**) Structural similarity network of predicted Dcr proteins, clustered based on TM-score ≥ 0.9. Each node represents a protein structure; edges indicate pairwise structural similarity above the threshold. Node colors match the clade classification defined in Fig. 2A (e.g., As1, As2, Ba1–Ba3, Mu1–Mu2), allowing direct comparison between structural and phylogenetic groupings. The most connected node (hub) from selected modules is shown with a red circle and its corresponding AlphaFold-predicted 3D structure is represented on the side. Proteins displayed correspond to representative members of non-canonical Dcr groups B, C, and E. Structural domains are colored as follows: PAZ (yellow), Helicase (green), RNase III (dark red/burgundy), DUF283 (cyan), and dsRBD (purple). Regions without domain predictions are shown in grey. Triangular nodes correspond to Dcr proteins with a canonical PAZ domain (cPAZ); oval nodes indicate non-canonical PAZ domains. (**c**) Phylogenetic tree of all Dcr proteins, with tips colored according to structural cluster membership, illustrating the correspondence between structural similarity and evolutionary lineage. “Other” corresponds to cluster 5-26.

The four major structural clades aligned well with the phylogenetic clades identified in our sequence-based analysis (Fig. 3C), indicating that despite extensive sequence divergence and domain variability, fungal Dcrs retain conserved structural architectures within evolutionary lineages. Cluster 1 contains sequences from clades As1 and Ba1, cluster 2 from As2, and cluster 3 from Ba2 and Mu1. Cluster 4 primarily contains Ba3, indicating that this is a unique lineage in Basidiomycota. Importantly, while proteins annotated as Dcr2 in UniProtKB majorly fall into Cluster 1, proteins annotated as Dcr1 are located into clusters 2, 3 and 4, indicating greater structural and likely functional diversity among Dcr1-labeled proteins. These findings underscore the need for systematic re-evaluation of Dcr annotations grounded in structural and evolutionary evidence rather than existing paralog labels.

The other cluster proteins (those falling outside the four major structural groups) may represent highly divergent Dcr variants, truncated sequences, or lineage-specific proteins with distinct structural features that deviate from the dominant structural folds. This group contains members from several phylogenetic clades, particularly dikarya fungi, and also includes Mucoromycota and Zoopagomycota Dcrs from clade Mu2 (Fig. 2B, 3C). It also includes all Dcrs from Blastocladiomycota clade Bl1, Miscrosporidia clade Mi1 and nearly all from Chytridiomycota clade Ch1, indicating substantial structural divergence in Dcr proteins in these non-Dikarya fungal lineages.

To further explore folding patterns, we examined the structure of the most connected (central) node within each major structural cluster (Fig. 3B). These representative proteins span different Dcr categories, with cluster 1 represented by a non-canonical D protein from *Aureobasidium namibiae* (clade As1), cluster 2 represented by *Li*Dcl1, a non-canonical E protein from *Lobaria immixta* (clade As2), cluster 3 represented by a non-canonical C protein from *Claroideoglomus candidum* (clade Mu1), and cluster 4 represented by *Sc*Dcl1, a non-canonical D protein from *Schizophyllum commune* (clade Ba3). Despite differences in domain composition detected by HMMER analysis, AlphaFold models of these proteins show broadly similar L-shaped Dcr architectures, comprising head, core and base regions. Importantly, the structural models revealed PAZ-like folds in the head region of representatives from clusters 1,2 and 4, even though cPAZ domains were not detectable at the sequence level in these proteins. This suggests that the RNA-binding functionality associated with PAZ might be structurally preserved in Dcrs despite substantial divergence, particularly in the more widely distributed non-canonical Dcr types.

### Ancestral state reconstruction supports an early origin and selective retention of the cPAZ domain in Fungi

To gain deeper insights into the evolutionary history of the PAZ domain in fungi, we performed an ancestral character estimation (ACE) analysis to determine whether the cPAZ domain was present in the last common ancestor of fungal Dcrs (Fig. 4A). The ACE analysis was based on the presence or absence of the cPAZ domain, mapped onto the Dcr phylogeny. The reconstruction suggests that the common ancestor of fungal Dcrs most likely harbored a cPAZ domain, supporting the hypothesis that this domain is an ancestral feature of fungal Dcrs. Despite its ancestral origin, the phylogenetic distribution of cPAZ shows strong lineage-specific conservation patterns (Fig. 4A), consistent with those observed in the phylogenetic analysis (Fig. 2D) and structural clustering (Fig. 3A, C). The presence of the cPAZ domain in these clades may reflect convergent retention under shared selective pressures, or alternatively, could be the result of horizontal gene transfer (HGT) from lineages such as plants or animals, where the cPAZ domain is nearly ubiquitous. Indeed, nearly all Dcr proteins in metazoans (99.9% of Dcrs) and plants (100% of Dcrs) contain a cPAZ domain in contrast to only 12.4% of fungal Dcrs (Fig. 4B). This stark difference likely reflects a greater evolutionary plasticity of the PAZ domain in fungi, where structural conservation can be retained despite extensive sequence divergence or domain reconfiguration. We evaluated the possibility of Dcr HGT from plants or animals to fungi, however our analysis did not find evidence supporting these events (Fig. 4C). These findings support a model in which the cPAZ was selectively retained in specific fungal lineages, while its apparent absence in most fungal Dcrs is most likely due to widespread sequence divergence rather than true domain loss. This interpretation is supported by the similar protein lengths observed between Dcrs with and without a cPAZ (Fig. 1D) and by the structural predictions showing a putative conserved PAZ-like fold in proteins lacking a detectable cPAZ (Fig. 3C).

**Figure 4.**
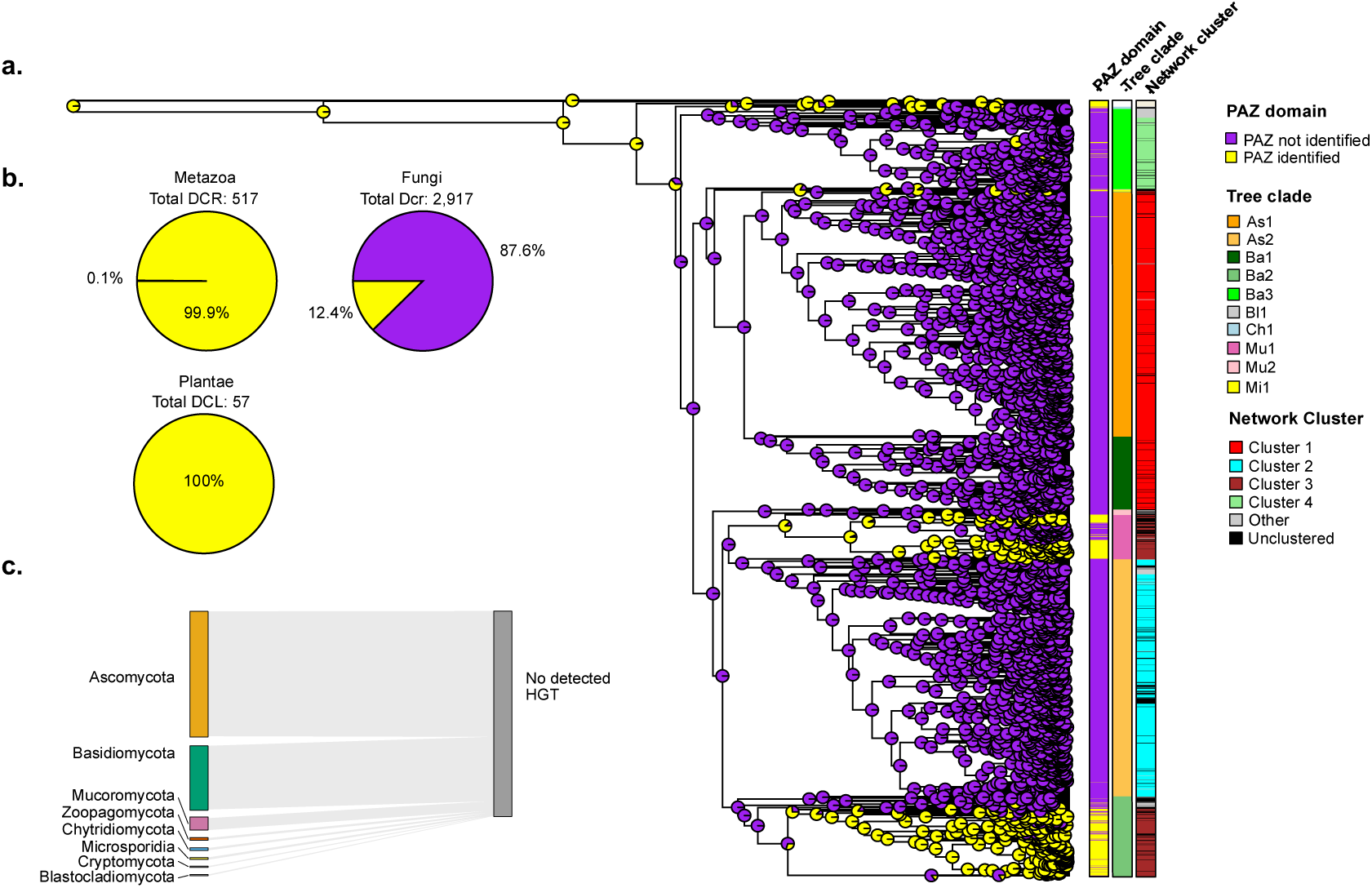
Evolutionary reconstruction of the PAZ domain in fungal Dcr proteins. (**a**) Phylogenetic tree of fungal Dcr proteins with ancestral state reconstruction for the canonical PAZ domain. Pie charts at internal nodes represent the inferred probability of PAZ domain presence (yellow) or absence (purple) using maximum likelihood estimation. Additional metadata layers include the presence/absence of a canonical PAZ domain, phylogenetic group (e.g., As1, Ba2, Mu1), and structural cluster as defined by the 3D similarity network in Fig. 3. (**b**) Proportion of Dcr proteins with a canonical PAZ domain across four eukaryotic kingdoms, Animalia, Fungi, and Plantae, highlighting variation in PAZ domain conservation. (**c**) Horizontal gene transfer (HGT) analysis. No inter-kingdom HGT events involving Dcr proteins were detected between fungi and plants, animals, or protists, indicating that the PAZ domain in fungal Dcrs evolved independently from those in other eukaryotic lineages.

### A sequence-divergent yet structurally conserved PAZ domain defines fungal Dcr diversity

We explored the possibility of a diverged fungal PAZ (fPAZ) in Dcr proteins lacking a detectable cPAZ. To address this idea, we compared the sequence and structural features of the cPAZ, present in canonical and non-canonical A and C Dcrs, with those of the putative fPAZ, which should be found in non-canonical B, D and E proteins. In proteins without a cPAZ, the fPAZ was defined as the region between the DUF and RNAse III domains. Using MEME (46), we identified four conserved motifs shared across both cPAZ and fPAZ domains. These motifs, designated Logo-1 through Logo-4, represent conserved sequence elements ordered from N-to C-terminal within the PAZ region (Fig. 5A). The logos revealed moderate to high positional conservation of aminoacidic residues across the motifs. Logo-1 displays highly conserved arginine (R) residues, and a conserved phenylalanine (F) at position 8. These residues reflect the characteristic composition of the 3’-end binding pockets in PAZ domains, where aromatic residues enable base stacking, and positive residues contribute to electrostatic interactions with the RNA(19, 47). Logo-2, similar to Logo-1, contains conserved F and tyrosine (Y) residues, followed by a stretch of basic residues, further supporting a conserved RNA-binding role. Logo-3 features several polar and charged residues, including K, D, N and Q, which are likely involved in maintaining the OB-fold structure of the PAZ domain via hydrogen bonding and overall stabilization. The presence of conserved K and N residues in the central region may also contribute to interactions with the phosphate in the RNA. Logos 1 to 3 were strongly associated with the cPAZ, being identified in over 80% of cPAZ-containing proteins, but in fewer than 20% of the sequences containing only an fPAZ. In contrast, Logo-4 was more broadly conserved across fungal Dcrs. It was present in 99.67% of the cPAZ sequences and in 56.71% of the fPAZ sequences. This motif is enriched in hydrophobic amino acids (F, I, P, A, L, M), consistent with a helical or turn-forming structure. Given its composition, Logo-4 likely localizes to the connector helix that links the PAZ to the first RNAse III domain, a region well-known to be essential for maintaining the spatial configuration required for precise RNA cleavage by Dcr (48). Mapping of these four motifs onto the three-dimensional structure of a representative canonical PAZ domain confirmed their spatial clustering at the RNA-binding interface (Supplementary Fig. 2). In particular, the motifs localize around the region that interact with the 3′ end of the dsRNA, supporting their proposed role in RNA anchoring and stabilization. The spatial proximity of Logo-1 and Logo-2 to the RNA entry groove aligns with their enrichment in basic and aromatic residues, while Logo-4 occupies a linker region adjacent to the RNase III domain, consistent with a role in domain coordination and structural integrity. This structural mapping provides further support for the functional relevance of these motifs in mediating RNA binding across diverse fungal Dcrs.

**Figure 5.**
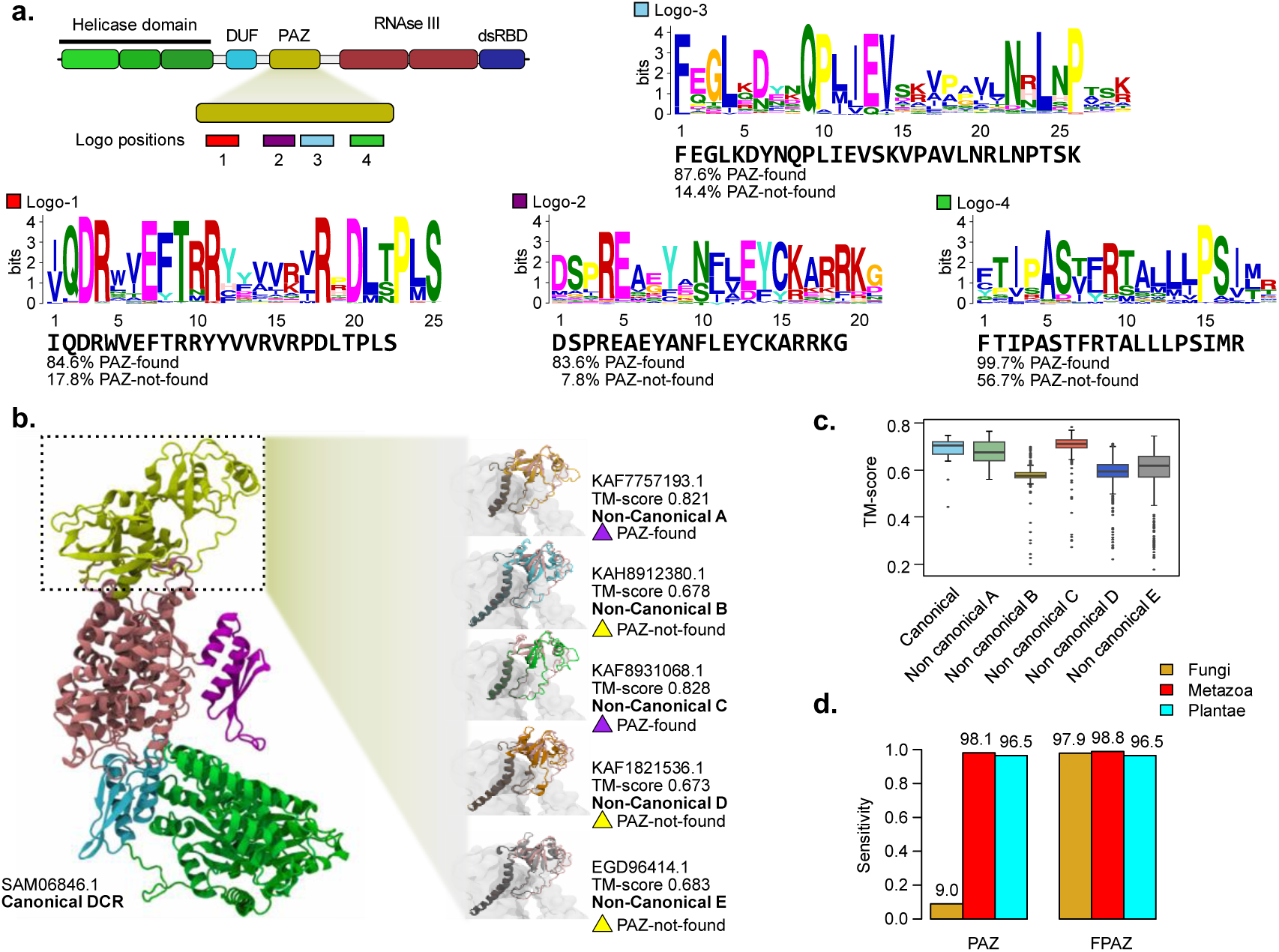
Motif conservation and structural variability of PAZ domains in fungal Dicer-like proteins. (**a**) Conserved sequence motifs identified within the canonical PAZ domain using MEME Suite. Schematic representation (top) shows the approximate location of four conserved motifs (Logos 1–4) along the PAZ region. Sequence logos illustrate consensus motifs and their relative frequency in proteins where the PAZ domain was detected (PAZ-found) versus those where it was not (PAZ-not-found). (**b**) Structural comparison of PAZ domains across Dcr categories. The full-length AlphaFold model of a canonical Dcr (SAM06846.1) is shown on the left, with the PAZ domain highlighted in yellow. Right panel: Representative AlphaFold structural models of PAZ domains from non-canonical categories A–E, annotated with TM-score values and presence/absence of a recognizable cPAZ domain. PAZ regions in non-canonical Dcrs are color-coded according to their category (A–E), as shown in the figure. Structural alignment was performed using the PAZ domain of SAM06846.1 as reference to comparing the spatial conservation of PAZ-like regions across non-canonical Dcrs. (**c**) Boxplot of TM-scores comparing structural similarity of PAZ domains across canonical and non-canonical Dcr categories. Canonical domains show consistently higher structural conservation. (**d**) Sensitivity comparison of the canonical PAZ HMM (PAZ) versus the fungal-specific FPAZ-HMM (FPAZ) across eukaryotic groups.

Next, we assessed the structural similarity of the cPAZ and divergent fPAZ domains. As a reference, we used the cPAZ domain from a representative canonical Dcr (Dcl1 from *Absidia glauca*). Structural comparisons revealed a high degree of similarity between this reference and the cPAZ domains of non-canonical Dcrs A (Dcl1 from *Entomophthora muscae,* TM-score of 0.821) and C (Dcl1 from *Dissophora ornata,* TM-score of 0.828). Although lower, structural similarity scores between the reference cPAZ and the putative fPAZ domains found in non-canonical Dcrs B, D and E (TM-scores of 0.678, 0.673 and 0.683, respectively), still indicate substantial structural conservation. This pattern of conserved fold architecture extends across the broader set of structural models analyzed (Fig. 5C).

Considering the high sequence divergence of fungal PAZ domains, and the under-representation of fungal Dcrs in the current PAZ HMM model, we generated a custom HMM-profile (FPAZ-HMM) based on a multiple sequence alignment of both cPAZ and structurally inferred fPAZ domains from our dataset. We used this profile to search for PAZ domains in Dcr-annotated proteins in UniProtKB. Compared to the current PAZ HMM, the FPAZ-HMM profile demonstrates markedly improved sensitivity for detecting PAZ domains in fungal Dcrs, while maintaining comparable performance in Metazoa and Plantae (Fig. 5D). Importantly, FPAZ-HMM was specifically trained to recognize PAZ domains associated with Dicer proteins-only, as other PAZ-containing proteins (PIWI, AGO) were intentionally excluded from the alignment used for model construction. Testing 236 proteins named AGO or ARGONAUTE found in the NCBI, we identified 0 instances of the FPAZ-HMM domain, suggesting high precision. In contrast, current general PAZ HMMs typically include PAZ domains from both Dcr and AGO proteins. These results support using of FPAZ-HMM as a robust and lineage-inclusive tool for detecting Dcr PAZ domains across eukaryotes and underscores the extensive sequence variability of PAZ domains across fungal clades.

### Structural and functional evidence support conserved RNA-binding activity of fungal PAZ domains

To gain insights into the functionality of the canonical and divergent fungal PAZ domains, we analyzed structural models of full-length canonical and non-canonical Dcrs predicted in complex with a pre-cleaved dsRNA substrate that was obtained from the PDB model of *D. melanogaster* Dicer (pdb_00007w0f) (Figure 6A, B). In these models, the dsRNA (green) interacts with the Dcr surface (grey), with key RNA-interacting residues highlighted in blue. In Dcrs with cPAZ, such as those from *Puccinia striiformis, Cryptococcus neoformans* and *Absidia glauca*, the PAZ domain and surrounding regions form a well-defined RNA-binding pocket. Basic residues, particularly K and R were consistently positioned within 3 Å of the dsRNA, forming a positively charged cleft that anchors the 3’ end of the RNA duplex (Fig. 6A, right). This configuration mirrors previously described PAZ-RNA interactions in plant and animal systems.

**Figure 6.**
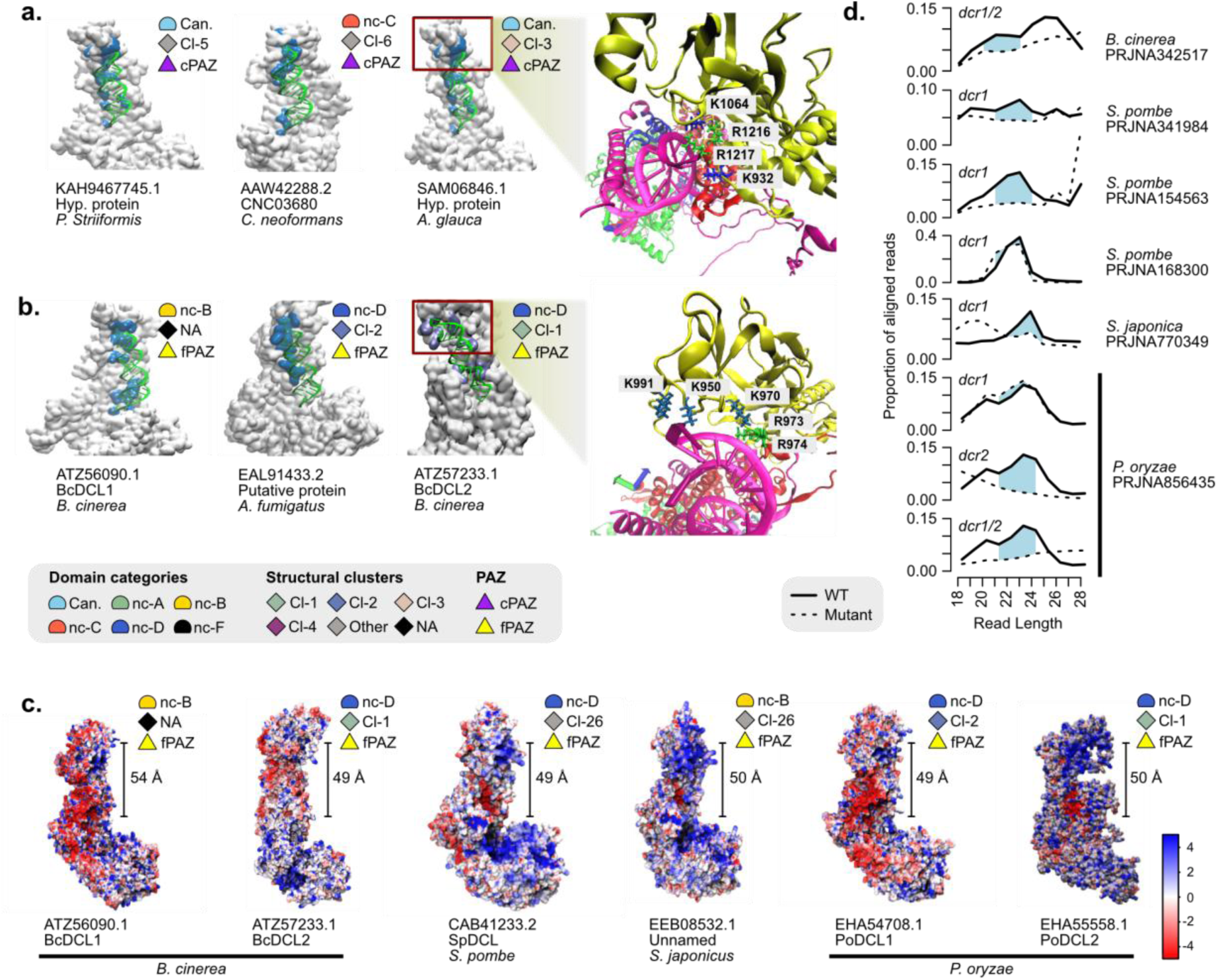
Structural and functional characterization of RNA-binding interfaces in canonical and non-canonical fungal Dcr proteins. (**a–b**) Surface representations of models of full-length Dcr proteins (gray) in complex with dsRNA (green), showing the RNA-binding interfaces (blue). (**a**) Canonical Dcr proteins containing a canonical PAZ domain (cPAZ), grouped by structural group (Cl-3, Cl-5, and Cl-6) and domain architecture. (**b**) Non-canonical Dcrs possessing fungal-specific PAZ regions (fPAZ), grouped by structural group (CI1, CI2, Other, or NA). Representative residues involved in interaction with RNA (lysines and arginines) are shown in atomic detail, with the side chains located within 3 Å of the RNA. Red boxes highlight the cPAZ or fPAZ region that anchors the 3’ end of the dsRNA. "Other" includes proteins belonging to minor structural clusters outside the four major groups. "NA" includes proteins that did not meet the clustering thresholds (≥90% TM-score similarity and pLDDT ≥70). (**c**) Electrostatic surface potential is shown for each model, with red indicating negative charge and blue indicating positive charge (scale in kT/e). The measured distance between the fPAZ region and the RNase III catalytic site (in Å) is shown, suggesting its possible role in determining the length of small RNA (sRNA). (**d**) sRNA library. Solid lines represent WT, dashed lines correspond to Dcr mutant. Light blue peaks indicate sRNA species lost in the mutant background.

Remarkably, similar interaction interfaces were observed in non-canonical Dcrs lacking a canonical PAZ domain (cPAZ), but containing a structurally defined fungal-specific PAZ (fPAZ), including Dcl1 and Dcl2 from *Botrytis cinerea* and the Dcr from *Aspergillus fumigatus*. These proteins display conserved K and R residues positioned in close proximity to the RNA substrate, supporting the idea that key electrostatic and structural features necessary for RNA anchoring are retained, even in the absence of canonical sequence motifs. To assess the conformational stability of these interactions, we conducted molecular dynamics simulations on a representative set of Dcr–RNA complexes. Across all variants analyzed—including canonical, non-canonical C, and fPAZ-containing non-canonical B and D— RMSD trajectories stabilized within the first 10–20 nanoseconds, indicating that the RNA-binding interface, particularly the PAZ or PAZ-like domain, maintains structural integrity throughout the 50-nanosecond simulations (Supplementary Fig. 3). These results support the functional relevance of the fPAZ fold as a stable RNA-interacting module across divergent fungal Dcrs.

To further explore the functional potential of the fungal PAZ regions, we analyzed the electrostatic surface distribution of representative Dcr proteins for which a function in sRNA biogenesis has been evaluated using mutants and sRNA sequencing. These include Dcl1 and Dcl2 from *B. cinerea*, Dcr1 from *S. pombe*, Dcr1 from *S. japonicus*, and Dcl1 and Dcl2 from *P. oryzae* (Fig. 6C). In these non-canonical Dcrs, the dsRNA-binding pocket displayed a predominance of positively charged residues, consistent with a role in RNA anchoring (Fig. 6C). Additionally, we measured the distance between the fPAZ domain and the RNAse III catalytic center, finding it to be approximately 50-55 Å, comparable to that reported in human Dicer (10). These structural configurations support the model in which PAZ domains act as a molecular ruler, securing the 3’end of the RNA and enabling precise cleavage by the RNAse III domains.

Finally, we compared sRNA length profiles from wild-type and mutant strains with disrupted dcr genes (Fig. 6D). In all cases examined, dcr mutation resulted in clear losses of distinct sRNA size classes, as indicated by the disappearance of specific peaks. These results confirm that non-canonical Dcrs contribute to sRNA biogenesis and strongly suggest that the fPAZ domain plays a critical role in RNA processing, despite its sequence divergence from canonical PAZ domains.

## Discussion

Understanding the structural diversity of Dcr proteins in fungi is essential. These proteins are central components of RNAi, a mechanism critical for gene regulation, genome stability, defense, and inter-species interactions. The dataset presented in this study represents, to our knowledge, the most extensive survey of fungal proteomes to date derived from annotated and reference-quality genomes, comprising 1,593 proteomes from 1,427 species across eight phyla and 44 taxonomic classes. While thousands of fungal genomes are publicly available, many are incomplete or lack standardized annotations, which limits their utility for comparative analyses of protein-coding genes such as Dcrs. Notably, our sampling includes a broader representation of non-Dikarya fungi compared to previous studies (21, 28), particularly within Mucoromycota and Microsporidia. This improvement reflects the increasing contribution of large-scale sequencing efforts such as the Joint Genome Institute’s 1000 Fungal Genomes Project (https://mycocosm.jgi.doe.gov/mycocosm/home/1000-fungal-genomes). Despite this broader coverage, several phyla are still poorly represented in our dataset, including Zoopagomycota (20 proteomes), Chytridiomycota (19 proteomes), Blastocladiomycota (2 proteomes), and Olpidiomycota (1 proteome), while other phyla are entirely absent. These limitations highlight the continued need to incorporate non-Dikarya and ecologically diverse fungal lineages into genome sequencing initiatives, to enable a more complete understanding of fungal molecular diversity.

Our survey of fungal Dcrs identified proteins broadly across fungal clades in the dataset, confirming that this machinery is widely retained across fungi, as others have found (21). Nevertheless, we found a relevant part of fungal species (∼10%) with no Dcr identified in their proteomes, according to our pipeline. While the loss of Dcrs is well established in budding yeasts within the Saccharomycotina and some Basidiomycota lineages (22, 49), broader surveys have shown that such losses may also occur sporadically across the fungal kingdom (31, 50). However, the extent of Dcr loss in other fungal groups remains less clear, largely due to the underrepresentation of non-Dikarya fungi in genomic datasets. In our analysis, most proteomes lacking detectable Dcrs belonged to Saccharomycotina, consistent with previous reports. However, we also identified apparent Dcr losses in non-Dikarya phyla. While these absences require further validation, they raise the possibility that RNAi pathways in fungi have evolved alternative mechanisms that operate independently of Dcr proteins. This contrasts with animal and plant systems, were the loss of Dcr typically results in severe developmental defects or lethality (51, 52) and suggests that fungi might possess a more flexible RNA silencing machinery. Consistent with this idea, evidence of Dcr-independent sRNA-processing by other members of the RNAseIII family has been reported in fungi, as is the case of MRPL3 in *Neurospora crassa,* or R3B2 in *Mucor lusitanicus* (45, 53). Fungal genomes encode a diverse array of RNases III which could play conceivable roles in RNAi. For example, our in-depth analysis of RNAse III-containing proteins uncovered a small number of potential DROSHA-like candidates. However, the general absence of the characteristic domain architecture typically associated with DROSHA suggests that these proteins may be highly divergent and thus require more targeted, in-depth analyses for definitive identification.

We found four major Dcr variants across fungi, but little is known about what the functional characteristics of these groups are. Among Dikarya, Dcrs from multiple clusters are found in organisms. This could relate to the radiation of RNAi functions in these organisms, as they contain a diverse complement of Dcrs. This diversity may reflect evolutionary pressure to fine-tune RNAi responses to different developmental or ecological contexts. In particular, the presence of sequence-diverging PAZ domains seems to be widespread, at least in the species represented in our dataset, with cPAZ domains limited number of species, mostly within Mucoromycota and Basidiomycota. Considering that ACE analysis supports that cPAZ was likely present in the last common ancestor of fungal Dcrs, this limited representation implies lineage-specific retention, which could be attributed to selective pressures to maintain a more precise or processive sRNA cleavage in these lineages. Both Mucoromycota and Basidiomycota comprise species engaged in complex ecological interactions, including mutualistic symbioses, such as Mycorrhizal associations, and pathogenic lifestyles. In these contexts, sRNA-mediated communication between fungi and their plant hosts may play a pivotal role in modulating host responses and facilitating successful colonization.

Interestingly, our phylogenetic analysis revealed that Dcr naming in databases have not been consistently applied across fungal lineages. Proteins carrying common names (Dcr1/Dcl1 or Dcr2/Dcl2) often appear in different clades, indicating that these labels do not accurately reflect true evolutionary or functional relationships. For example, proteins annotated as Dcr1/Dcl1 are distributed across Ba2 and Ba3 in Basidiomycota, which form separate phylogenetic and structural clusters. Similar patterns are observed for other Dcr annotations. These observations highlight that current Dcr naming conventions are largely disconnected from the structural and phylogenetic diversity of fungal Dcr proteins, underscoring the need for a more systematic, structure- and phylogeny-informed classification framework.

In fungi, the PAZ domain has traditionally been considered dispensable for Dcr function, since various well-characterized fungal models, including *Schizosaccharomyces pombe, Neurospora crassa*, *Trichoderma atroviride*, *Aspergillus fumigatus*, or *Botrytis cinerea,* harbor non-canonical Dcrs that lack detectable PAZ domains but remain functional in sRNA production (22, 40–45). However, our structural analyses challenge this notion. Many of these non-canonical Dcrs retain a PAZ-like fold capable of anchoring dsRNA despite lacking sequence similarity to canonical PAZ domains. This finding suggests that structural conservation, rather than primary sequence similarity, may underline the preservation of PAZ functionality in fungi.

Detailed structural comparisons revealed strong conservation of R and K residues in the binding pocket pointing to a conservation of the dsRNA-protein interface. However, we found that the highly hydrophobic Logo-4 motif is much more conserved in fungal Dcrs. This hints that the overall structure of the PAZ domain is more conserved among fungi, while the actual binding interface could be more lineage specific (Logos-1, -2, -3). Consistent with previous studies (54), we found that PAZ-like domains in fungal Dcrs conserve the R and K residues critical for 3′ end binding. Amino acids like glutamic acid (E), R, and K are often involved in ionic interactions and hydrogen bonding, essential for dsRNA binding. Y and F contribute to hydrophobic interactions, stabilizing the RNA-protein complex. The identification of these conserved residues and their role in ionic interactions, hydrogen bonding, and hydrophobic stabilization strengthens the hypothesis that even non-canonical PAZ domains can perform essential functions. Understanding these conserved motifs not only shed light on the molecular mechanisms by which fungal Dcrs interact with dsRNA but also opens avenues for functional validation and targeted manipulation. For instance, residues such as R and K, which mediate 3′ end binding, as well as other amino acids involved in electrostatic and hydrophobic interactions, are promising candidates for site-directed mutagenesis to assess their specific contribution to RNA binding and cleavage efficiency. These experiments would help clarify whether structurally conserved PAZ-like folds can fully compensate for the absence of canonical PAZ domains in non-canonical Dcrs. Moreover, identifying lineage-specific conservation patterns, such as the highly preserved hydrophobic Logo-4 motif, could inform the design of small molecule inhibitors or RNA mimetics that selectively disrupt dsRNA recognition in certain fungal groups. This may have practical implications for biotechnological or agricultural applications, such as developing of fungal RNAi-based tools for gene silencing or plant-pathogen interaction control. Ultimately, resolving how these motifs contribute to RNA processing in structurally divergent Dcrs will enhance our understanding of fungal RNAi flexibility and evolutionary innovation.

Our findings reveal that fungal Dcr proteins maintain essential RNA-binding function through structurally conserved domains, even when canonical sequence motifs are undetectable. This observation challenges the central assumption in many genomic annotations, that domain absence inferred from sequence similarity implies functional loss. In fact, we show that structural and electrostatic features can persist despite extensive sequence divergence, preserving the biochemical properties of RNA-binding interfaces.

The difficulty of sequence-based domain models (e.g., InterProScan, Pfam HMMs) to detect PAZ-like regions in fungal Dcrs exemplifies this limitation. These tools rely on hidden Markov models trained primarily on canonical sequences from animals and plants and may fail to detect highly divergent variants that preserve structural and functional features despite lacking recognizable sequence similarity. This issue has also reported in recent work using structure-based HMMs (55). As a result, structurally conserved domains can remain undetected, leading to underestimation of functional conservation across divergent eukaryotic lineages (33).

Although our primary goal was to improve domain detection in fungal Dcrs, FPAZ-HMM also demonstrated robust performance in Dicer proteins from plants and metazoans available in NCBI. Comparative evaluation against the canonical PAZ model (PF02170) showed high sensitivity for known Dcrs, while avoiding annotation of PAZ in AGO proteins. This precision and sensitivity across diverse fungi suggests this may be valuable to search for Dcr in other less-studied eukaryotic lineages, such as protists, microalgae, or eukaryotic parasites more broadly. Such taxa often exhibit high sequence divergence and are underrepresented in canonical domain models, a challenge previously described in other divergent lineages using structure-informed HMMs (56, 57). Our strategy demonstrates that structurally informed HMM profiles which include more diverse proteins can improve annotation quality in such groups, uncovering hidden domain conservation and guiding functional predictions in underexplored branches of the eukaryotic tree. Overall, this highlights FPAZ-HMM as a useful complementary profile to canonical PAZ.

Beyond the methodological implications, fungal Dcrs emerge as a compelling model for understanding modular protein evolution, where domains can be lost, diverged, or structurally reconfigured without abolishing function. This plasticity likely reflects selective pressures to streamline or adapt RNAi responses to diverse ecological contexts, pressures that may have also shaped other multidomain regulatory proteins across eukaryotes. By combining phylogenetic analysis, structural modeling, and motif detection, our work highlights the power of integrative approaches to uncover hidden functional conservation in rapidly evolving systems.

In addition to evolutionary insights, the recognition of structurally conserved RNA-binding domains in non-canonical fungal Dcrs has practical implications. Several fungal species carrying such divergent Dcrs, including *Botrytis cinerea*, *Trichoderma atroviride*, and *Puccinia striiformis*, are agriculturally relevant pathogens or symbionts. Understanding how their RNAi machinery tolerates domain loss could inform the design of fungal-specific gene silencing tools, including minimal RNAi constructs for functional genomics, or novel strategies for crop protection via cross-kingdom RNAi.

## Materials and Methods

### Recovery and quality assessment of fungal proteomes

We retrieved a total of 1,593 annotated and reference fungal proteomes from the NCBI (data available up to November 2022), corresponding to 1,427 unique species and 172 duplicated species from different strains. We used the the Entrez esearch tool via its command line interface (Kans, 2013), with the query ’"Fungi"[Organism] OR fungi[All Fields]) AND (fungi[filter] AND "representative genome"[filter] AND (all[filter] NOT anomalous[filter] AND all[filter] NOT partial[filter]) AND "has annotation"[Properties]’, applying filters to select only representative genomes with available annotations, while excluding anomalous or partial assemblies.

We used BUSCO (Benchmarking Universal Single-Copy Ortholog assessment) (58) v.5.3.2 to evaluate the completeness of proteomes. We first applied the universal_odb10 and fungi_odb10 dataset, followed by additional phylum-specific datasets when available, including asomcyota_odb10, basidiomycota_odb10, microsporidia_odb10, and mucoromycota_odb10. We retained all retrieved proteomes for posterior analyses, regardless of their completeness score in BUSCO. We made this decision to maximize the detection of Dcr proteins, ensuring that potentially relevant sequences from less complete proteomes were not excluded.

### Recovery of profile Hidden Markov Model and identification of putative Dcr proteins

We retrieved a set of profile Hidden Markov Models (HMMs) representing highly conserved Dcr domains within eukaryotic species from the Interpro database (ebi.ac.uk/interpro). These HMMs have been used in previous Dcr studies (22, 36, 59) and include PF00270/IPR011545 (DEAD/DEAH box helicase subdomain), PF04851/IPR006935 (ResIII helicase subdomain), and PF00271/IPR001650 (Helicase C terminal helicase subdomain), PF00636/IPR000999 (RNase III domain), PF02170/IPR003100 (PAZ domain), PF03368/IPR005034 (DUF/Dicer dimerization domain), and PF00035/IPR014720 (double-stranded RNA binding domain). We scanned the fungal proteomes using the HMMER v.3.3.1 toolkit (60) to identify putative Dcr proteins. The thresholds for detection of a domain were set at an E-value ≤10^-3^, and only proteins containing at least one RNase III domain (PF00636/IPR000999) were retained for further analysis.

### InterProScan analysis

InterProScan v.5.44 (61) was used to scan protein sequences. All analyses were conducted using default parameters.

### In silico protein structure prediction

To predict the three-dimensional structures of the Dcr proteins, we used the local version of Colabfold (62) v1.5.5, a Google Colab implementation of Alphafold2 (35). In the first step, multiple sequence alignments (MSA) were generated against all available databases using the “DB load mode 3” setting, which preloads all sequence databases into memory to speed up alignment searches. We executed this process using the following command: colabfold_search --db-load-mode 3 --threads 54 dcr_sequences.fasta database a3m_output. Subsequently, we predicted the protein structures using the same software with the command: colabfold_batch --num-models 1 --num-relax 0 msas_input/ pdb_output/. To reduce computational time, we only generated one structural model per protein, and no structural relaxation was performed. We selected protein models with a predicted Local Distance Difference Test (pLDDT) score ≥70 for further analysis, ensuring the inclusion of structures with high predicted accuracy for downstream analyses.

### Electrostatic Surface Calculations

To investigate the electrostatic surface distribution of selected Dicer proteins, we applied a two-step computational workflow consisting of structure preparation and electrostatic potential calculation. Protein structure were preprocessed using PDB2PQR (63), which assigns atomic charges and radii according to the PARameters for Solvation Energy (PARSE) force field and determined protonation states at pH 7.0 using PROPKA (64, 65). This preprocessing ensures consistent treatment of ionizable residues, addition of hydrogens, and optimization of side-chain orientations for electrostatic calculations.

Electrostatic potentials were then computed using the Adaptive Poisson–Boltzmann Solver (APBS) (66). We performed the calculations using the linearized Poisson–Boltzmann equation (LPBE), with the solute and solvent dielectric constants set to 2.0 and 78.54, respectively. We used a solvent probe radius of 1.4 Å, temperature of 298.15 K, and ion-accessibility parameters (swin = 0.3 Å, sdens = 10.0). We set surface and charge discretization methods to smol and spl2, respectively. We set the boundary condition to sdh (single Debye– Hückel), and we automatically generated the electrostatic grid to encompass each protein’s structure with sufficient resolution. Electrostatic potential maps were output in OpenDX format and used to visualize surface charge distributions, with a focus on the double-stranded RNA binding regions. Total electrostatic energies were also computed but not directly analyzed.

### Generation of Protein Similarity Networks and Protein Structural Networks

We generated a sequence similarity network using the Sequence Similarity Network pipeline v.1.0.0 (https://github.com/MiguelMSandin/SSNetworks). In brief, this pipeline performs a local pairwise alignment of sequences using BLASTn to calculate similarities between all sequences in the dataset. We processed the resulting pairwise similarities to remove reciprocal hits and self-hits, ensuring a clean input for network construction. We generated the network considering an identity threshold of 50% and a coverage of 80%. For the protein structural network, we aligned the obtained protein structures in PDB format using mTM-align (67). We generated the structural network considering TM-scores of 90% or more between nodes. We used Cytoscape (68) v.3.10.0 to visualize and analyze the networks.

### Phylogenetic tree generation

We performed multiple sequence alignments using T-Coffee (69) v13.45.0 with default parameters. Proteins corresponding to each of the 149 clusters generated by the sequence similarity network analysis were aligned independently. Subsequently, we carried out profile alignment for the 149 alignments to generate a global alignment. For this, we used ClustalW (70) v.2.1 with the-profile option. The global protein alignments were trimmed using ClipKit (71) v2.1.1 with default parameters. We inferred the best-fit model and maximum-likelihood phylogenetic trees from the trimmed sequence alignments using IQ-TREE (72) v2.2.5 with the following parameters: -B 1000 -m MFP -nt AUTO.

### Ancestral reconstruction

We used the fitmk function of the phytools (73) package to fit a discrete Markov model for the ancestral reconstruction of the Dcr proteins, taking as input the phylogenetic tree generated above. In this case, we used the "Equal Rates" (ER) model, which assumes equal transition rates between states, with the following command: fitER = fitMk(midroot_tree, PAZ_level, model = "ER"). The anc.recon function was used based on the fitted model, and we performed ancestral reconstruction using the marginal method to infer the most probable states at the ancestral nodes of the tree, using the command: fit.marginal = ancr(fitER, type = "marginal").

### Horizontal gene transfer analysis

To assess the potential occurrence of horizontal gene transfer (HGT) among fungal Dcrs, we analyzed Dcr sequences using HGTector (74) v2.0b3. The search module of HGTector was executed with DIAMOND(75) against a predefined NR reference database (40,310 genomes, retrieved in January 2023). We run the ‘analyze’ command using default parameters, except for the --self parameter, which was set to 4751 (NCBI taxonomy ID for Fungi) to filter intra-fungal homologs.

### Docking and molecular simulations

We performed docking of the Dcr-RNA complex using HDOCK (47) v1.0. Protein models of Dcr, were provided in PDB format, and were used along with the dsRNA derived from the Drosophila Dicer2 PDB model (7W0F). We performed *ab initio* docking, defining the residues in the receptor (Dcr) and ligand (RNA) where preferential binding was expected.

For the molecular dynamics simulations, we used the CHARMM36 (76) force field to construct the protein structure file (PSF) for both protein and dsRNA. The simulation workflow consisted of three stages. First, energy minimization was performed using 20,000 steps to remove steric clashes and stabilize the initial system using a restriction of 1 kcal/mol. Next, an NVT equilibration phase was conducted for 0.25 ns constraints applied to the protein and dsRNA to maintain structural integrity during thermal adjustments (1 kcal/mol). Finally, production simulations were carried out in two phases: 50 ns simulations with dsRNA restrictions maintained, followed by an additional 50 ns simulations with all constraints released to allow full system flexibility. Each molecular simulation was performed in three independent replicates to assess the reproducibility and stability of the results. We generated all molecular dynamics steps using NAMD3 (77). The equations of motion were integrated with a 2-fs time step using the Verlet algorithm. Langevin dynamics (damping coefficient of 1 ps) and the Nosé–Hoover Langevin piston method was employed to maintain constant temperature (310 K) and pressure (1 atm). Long-range electrostatic interactions were calculated using the Particle Mesh Ewald (PME) method, and van der Waals forces were computed with a cutoff of 12 Å.

### Motif discovery in protein sequences

We performed motif discovery using the Multiple Em for Motif Elicitation (MEME) tool v. 5.5.7 from the MEME suite (46). We ran MEME locally with the options -protein -mod zoops-nmotifs 4 -minw 6 -maxw 50. We used the obtained motifs to scan the sequences with the Find Individual Motif Occurrences (FIMO) from the MEME suite. We ran FIMO with default parameters.

### Construction of a custom HMM profile for the PAZ domain

To generate a high-confidence HMM profile of the PAZ domain, we selected 2,452 PAZ-containing regions from fungal Dicer proteins, including both canonical (cPAZ) and non-canonical (fPAZ) representatives identified in our dataset. We first aligned sequences separately using MAFFT (78) v7.505 with the --auto option, visually inspecting alignment quality to confirm the presence of informative blocks. Poorly aligned or uninformative regions were trimmed using ClipKit v1.3.0 in kpi mode. We then merged the resulting canonical and non-canonical alignments via profile-profile alignment using ClustalW v2.1. This combined alignment was once again trimmed with ClipKit to remove potential residual noise.

We used the resulting filtered alignment to build a PAZ domain-specific HMM profile using hmmbuild from the HMMER suite v3.3. The final model consisted of 395 match states (effective sequence number = 24.25; relative entropy per position = 0.59), indicating moderate diversity and information content. Attempts to improve the entropy score using the--symfrac 0.7 option did not yield significant differences.

We evaluated the sensitivity (recall) of the model using hmmsearch against several curated datasets, including fungal Dicer proteins identified in our pipeline and representative Dicer-like proteins from plants and animals. True positives (TP) were defined as sequences successfully detected by the FPAZ-HMM model, and false negatives (FN) as Dicer sequences in the dataset that were not detected. Recall was calculated as TP / (TP + FN), following the conventional formula. Although we did not formally assess specificity, we verified that the model did not detect PAZ domains in a set of unrelated negative controls, including Argonaute proteins and Arabidopsis thaliana EIL1, supporting its discriminatory capacity.

### Processing of sRNA-seq libraries

We performed a comprehensive search of publicly available sRNA-seq datasets in NCBI, utilizing the NCBI-SRA tools, eDirect, and custom python scripts using fungal-related terms (e.g., fungi, fungus, specific fungal genera) combined with sRNA-related keywords (e.g., miRNA, microRNA, milRNA, small RNA*, sRNA, siRNA, smRNA, epigen*, RNAi). We cross-referenced the publications matching the search criteria with NCBI sample databases (SRA, Bioprojects, Biosamples, and GEO datasets) to identify relevant datasets. We manually reviewed the entries to identify BioProjects containing libraries likely including fungal sRNA-seq data from wild-type or Dicer mutants. We processes the selected libraries using the YASMA small RNA analysis suite (79) (github.com/nateyjay/yasma). We downloaded libraries using SRAtools. 3’ adapters were identified using YASMA-adapter and trimmed with cutadapt (Martin, 2011) with the following settings: cutadapt -a [adapter] --minimum-length 15 --maximum-length 50 -O 4 --max-n 0 --trimmed-only -o [out_file] [file]. We aligned trimmed sequences to the closest matching reference genome using YASMA-align, which mimics the unique-weighting approach found in ShortStack3 (80). sRNAs were classified into general sizes using YASMA-size-profile.

### Determination of molecular distances in protein structures

We used the Bio3D R library (81) to calculate the molecular distance between the N-terminus of the PAZ domain and the N-terminus of the RNAse IIIa domain. The protein structure, provided in PDB format, was imported to R using the read.pdb function and the coordinates of the alpha-carbon atoms corresponding to the N-terminal residues of the PAZ and RNAse IIIa domains were extracted. We calculated the Euclidean distance between these coordinates in Angstroms using the xyz.dist function in Bio3D.

### Data availability

Structural models of fungal Dicer-like (Dcr) proteins predicted by AlphaFold2, along with the custom Hidden Markov Model (FPAZ.HMM) used for PAZ domain identification, have been deposited in Figshare and are publicly accessible at: https://doi.org/10.6084/m9.figshare.28592981.v1.

All other data supporting the findings of this study are available from the corresponding authors upon reasonable request.

## Acknowledgements and funding sources

This work was funded by Agencia Nacional de Investigación y Desarrollo (ANID)-National Fund for Scientific and Technological Development (FONDECYT) Grants 11220727 (to N.R.J.), 11251927 (to C.M.), 1221667 (to V.C-F.), 3230600 (to P.V.), 11200209 (to J.P.C.), and 1221064 (to J.A.R-P); the Concurso Interuniversitario de Iniciación en Investigación Asociativa Grant IUP23-26 (to J.P.C.); ANID-Millennium Science Initiative Program, Millennium Institute for Integrative Biology iBio Grant ICN17_022 (to C.M., N.R.J., and E.A.V.); ANID-Millennium Science Initiative Program, Millennium Nucleus in Data Science for Plant Resilience Phytolearning Grant NCN2024_047 (to C.M. and E.A.V.); ANID-Vinculación Internacional Grant FOVI230159 (to E.A.V. and C.M.); ANID-Beca Doctoral 2021-21211564 (to B.V-V.) and 2022-21221055 (to I.O.); and Beca Doctoral Universidad Mayor (to L.M.). The authors acknowledge the use of the High-Performance Computing UOH Laboratory (FIC 40059065-0) at Universidad de O’Higgins, Rancagua, and the computational infrastructure of the Center for Genomics and Bioinformatics at Universidad Mayor, Chile, for providing supercomputing support.

